# Target deletion or holding on sections after enzyme digestion monitored with attenuation-of-sound images

**DOI:** 10.1101/2025.03.14.643196

**Authors:** Katsutoshi Miura, Toshihide Iwashita

## Abstract

Tissues consist of various components and if these can be deleted or reserved, their location and proportion can be detected. Scanning acoustic microscopy (SAM) calculates the attenuation of sound (AOS) through tissue sections to obtain histological images without staining. AOS values are reduced as tissue components break down. Here, we digested specific components in tissues using enzymes and followed the process with AOS imaging over time. In addition, we applied specific dyes and antibodies to inhibit enzyme activity and maintain a specific component in the section.

We used specific enzymes to degrade tissues that contain the enzyme’s substrate, such as collagenase for bone, elastase for skin and arteries, actinase for amyloid-positive cervical arteries and lymph nodes, amylase for the corpora amylacea (CA) of the brain and DNase and haematoxylin for adenocarcinomas.

Collagenase digested bone and cartilage to clearly visualise the internal structure. The structural components had characteristic AOS values, which gradually decreased. Elastases break elastic fibres in the skin and arteries differently between young and old individuals. The dermis and tunica media of arteries in the elderly fracture more easily than those of younger individuals. Actinase digested the cervical artery except for amyloid deposits, which were preferentially detected by Congo red staining. Actinase-digested lymphoid cells remained horseradish peroxidase (HRP)-staining positive. Amylase digested some CAs, which became periodic acid-Schiff (PAS) staining negative and diminished in size by electron microscopy observation. Cell nuclei were digested and deleted by DNase except for those stained with HRP. Residual nuclear images of AOS matched those of light microscopy, and haematoxylin staining inhibited DNase digestion of the nucleus.

Specific inhibition of enzymes preserved the target cells and materials. SAM observation can monitor the tissue breakdown process, which provides an advantage over light microscopy as no staining is required and exhibits higher sensitivity to detect fragile structures.

## Introduction

Scanning acoustic microscopy (SAM) detects tissue viscosity by measuring the attenuation of sound (AOS) [1–3]. Tissues consist of various components that maintain their mechanical strength and active function. If a particular component breaks down, the tissue structure and function are damaged, which can be used to identify the role and constituent. Enzymes that bind to specific substrates can break down a particular constituent for its detection. The more that a substrate is digested, the more easily ultrasound can pass through the structure, which reduces the AOS values. In this study, we digested various tissues with substrate-specific enzymes and followed their AOS images over time. Moreover, we used specific enzyme inhibitors to protect and preserve the target.

SAM constructs histological images by plotting the AOS values of each region in the section [4,5]. SAM uses the same section as that for light microscopy (LM), although its thickness may be twice or three times that of usual LM sections. This thick section contains many constituents and provides a large amount of substrate for the enzymes to digest. Enzyme activity can be monitored over time by AOS images because SAM observation does not require staining and takes a few minutes to generate the images.

In a previous study, we used various proteases to elucidate pathological alterations, such as actinase to detect amyloid deposits [2], collagenase to reveal the structural differences of the ageing lung [6], renal artery [7], aorta [8] and aortic valve [9] and pepsin to assess skin ageing [10].

In this study, we attempted to eliminate various tissue components, including proteins, glycans and nucleic acids, to clarify their location. Moreover, we used materials that block enzyme activity, such as dyes and antibodies, to preserve specific cells and materials.

This method allows for the intentional deletion or retention of components in a section and the degree of degradation can be compared using the AOS values.

## Materials and methods

### Human and mouse specimen preparation

Patient samples were retrospectively selected from the files of Hamamatsu University School of Medicine between 2008 and 2022. The following data were collected: age, gender, pathological diagnosis from 1/1/2008 to 31/12/2022. Paraffin sections and cytology slides without a link to the patient’s identity were used in this study. The study protocol conformed to the ethical guidelines of the Helsinki Declaration of 1975, as revised in 1983, and was approved by the ethical committee of the Hamamatsu University School of Medicine (approval no. 19-180). All procedures were conducted according to the guidelines and regulations of the ethical committee. A waiver of consent was obtained for each patient because of anonymized ordinary samples. After data collection, authors didn’t access to information that could identify individual persons. Tissue samples were fixed in a 10% buffered formalin solution, embedded in paraffin and sliced into flat sections (10-µm-thick sections were prepared for SAM, whereas 4-µm-thick sections were prepared for LM).

For mouse bone tissue, the bones were provided by Dr. Y. Enomoto from Department of Regenerative & Infectious Pathology and fixed and soaked in 0.5 mol/L ethylenediaminetetraacetic acid (EDTA) solution (Fujifilm Wako chemicals, Tokyo, Japan) for 2 days for decalcification.

For the cytology section, residual free cells from ascites or pleural effusions were prepared to make single-cell-layer slides using a previously reported liquid-based cytology method (BD CytoRich™; Franklin Lakes, NJ, USA) [11]. The slides were fixed in 95% ethanol for 10 min and postfixed in 10% buffered formalin for 45 min.

### Enzyme digestion

To digest the sections, various enzymes were used, including actinase E (pronase E) (Funakoshi, Tokyo, Japan), collagenase type 2 (Worthington, Lakewood, NJ, USA), elastase (Worthington), DNase 1 (Merck, Darmstadt, Germany) and α-amylase (Fujifilm Wako, Osaka, Japan). Actinase, collagenase type 2 and elastase were dissolved at 1 mg/mL in phosphate-buffered saline (PBS) containing 0.5 mM CaCl_2_. α-Amylase (10 mg/mL) was dissolved in PBS (pH 7.4) and DNase 1 (0.1 mg/mL) was dissolved in 20 mM HCl containing 1 mM MgCl_2_ and 1 mM CaCl_2_. The enzyme solution was mounted on the section and incubated at 37°C. The activity of each enzyme determined the incubation duration. The sections were washed in distilled water at each time point and observed with SAM. After observation, the same section was reincubated in the enzyme solution.

### SAM observations

We used a SAM system (AMS-50AI; Honda Electronics, Toyohashi, Aichi, Japan) with a central frequency of 320 MHz and a lateral resolution of 3.8 µm, as previously reported [12,13,14]. The tissue or cytology section was placed on the stage and distilled water was used as the coupling fluid between the transducer and the section. The waveforms reflected from the surface and bottom of the sample were compared to measure the AOS at each point [2]. The waveform from the glass surface without a specimen was considered the zero AOS area (black) and was used as the reference. The transducer scanned the sections along the X- and Y-axis for a few minutes to capture the images. The plotted AOS value at each point generated an AOS image.

### LM observation

LM slides taken from near the SAM section locations or the same section as for the SAM observation were prepared for comparison with the corresponding AOS images. Staining methods, including haematoxylin and eosin, Elastica Masson trichrome, Elastica, Congo red and periodic acid-Schiff (PAS), were the same as the routine histology methods employed in the pathology laboratory.

### Immunohistochemistry

We utilised the Dako REAL EnVision detection system using the peroxidase reaction with DAB for immunohistochemistry and followed the analysis procedure. The primary antibody was anti-Ki-67 (MIB-1, DAKO). For antigen retrieval, histological and cytological sections were soaked in 10 mM Tris-EDTA buffer (pH 9.0) (Abcam, Tokyo, Japan) at 95°C for 40 min. After immunostaining, the sections were counterstained with haematoxylin.

### Transmission electron microscopic observation of paraffin sections

Transmission electron microscopic (TEM) observation of formalin-fixed paraffin sections was performed using previously reported methods [15,16]. DAB-stained sections were fixed with 2% glutaraldehyde for 1 h for pre-fixation and then incubated with 2% osmium tetroxide for 15 min for post-fixation. The sections were dehydrated with an alcohol gradient and embedded in epoxy resin (Quetol 812, Nisshin EM Co.) by heating at 60°C for 48 h. Ultrathin 70-nm-thick sections were prepared, stained with lead and uranium acetate (Merck) and observed by JEM 1400 Plus (JEOL, Tokyo, Japan).

## Results

### Collagenase type 2 degradation of the mouse bone

Collagenase type 2 is an enzyme that degrades the collagen in cartilage and bone and after incubation, the AOS images showed a decreased value over time (Fig. 1). Before digestion, the tissue was a mixture of many components with no gaps between them, making it challenging to distinguish the boundaries between the components. The marrow was easy to discriminate from the trabeculae. Throughout digestion, the AOS values of all components decreased (Fig. 2). The AOS values of components in increasing order are the tendon, periosteum, trabeculae, marrow and cartilage and this order was maintained up to 60 min of digestion. Trabeculae dropped in rank because of decalcification treatment.

**Fig 1.**
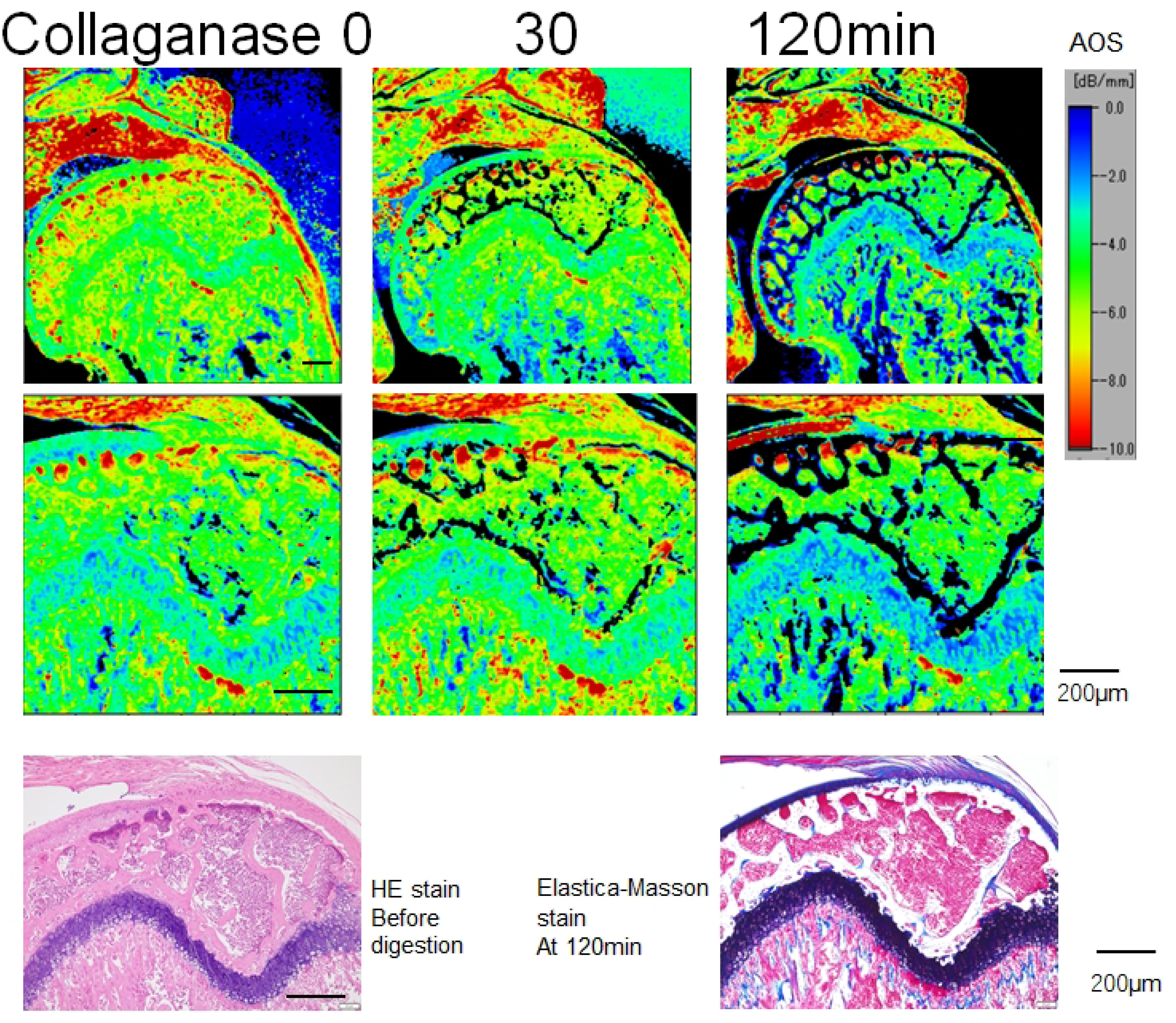
Collagenase type 2 degradation of the mouse bone. The femoral head of the mouse bone was digested using collagenase type 2. The figures in the upper rows show the attenuation-of-sound (AOS) images, which reveal a gradual decline. Cartilage and bone collagens were digested, thereby reducing the AOS values, and the bone marrow haematopoietic cells withstood the digestion. The ligaments on the articular cartilage and the connecting portions between the cartilage and trabeculae maintained high-AOS values.

**Fig 2.**
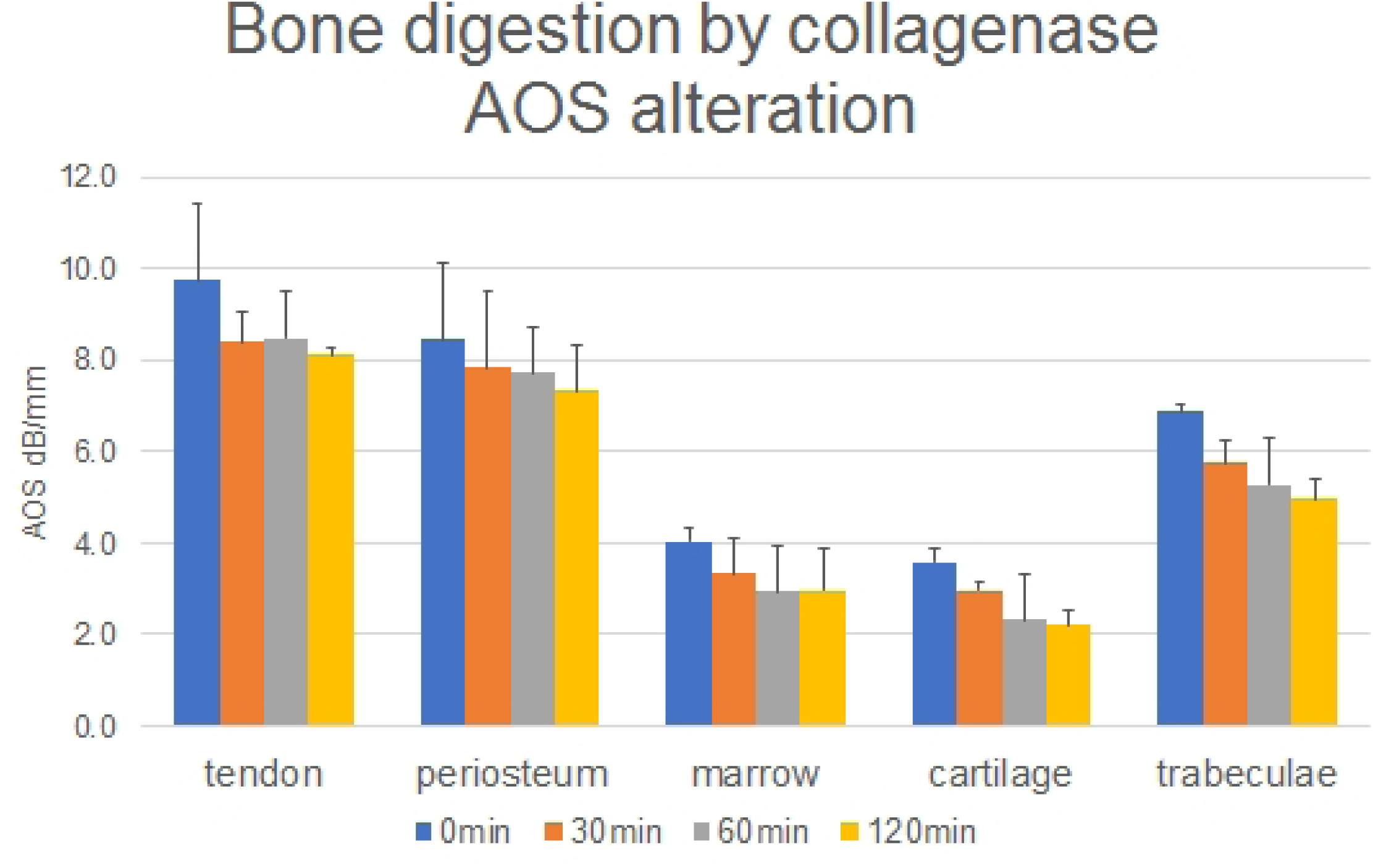
Mean and standard deviation of attenuation-of-sound (AOS) values after collagenase digestion. Collagenase digested the bone trabeculae, cartilage, tendon, periosteum and bone marrow tissue and their AOS values were obtained at 30, 60 and 120 min after digestion. From the beginning, the bone components consisted of high-AOS and low-AOS values. The high-AOS value group included the bone trabeculae, tendons and periosteum and the lower AOS value group consisted of cartilage and bone marrow. The AOS values of all components gradually decreased. N = 3 for each component.

### Elastase degradation of the facial skin of elderly and young individuals

The skin of the elderly is characterised by a thin epidermis, flat epithelial rete ridges, thin dermis and a few adnexal glands. On the other hand, the skin of the young is distinguished by a thick epidermis, elongated epithelial rete ridges, a dense dermis and many adnexal glands. In the elderly dermis, elastic fibres are clumped together under the epidermis as solar elastosis. In younger individuals, the elastic fibres are spread evenly among the thick collagen fibres of the dermis.

The upper row of Fig. 3 shows AOS images of the facial skin of 3- and 79-year-old patients. In the young skin, the epidermis decreased in AOS values, except for the surface keratin layer and the dermis maintained high-AOS values over time. In the elderly skin, the epidermis exhibited reduced AOS values, except for the surface keratin and basement membrane and partial portions remained unchanged. The AOS values of the dermis were reduced, especially in the papillary region corresponding to solar elastosis.

**Fig 3.**
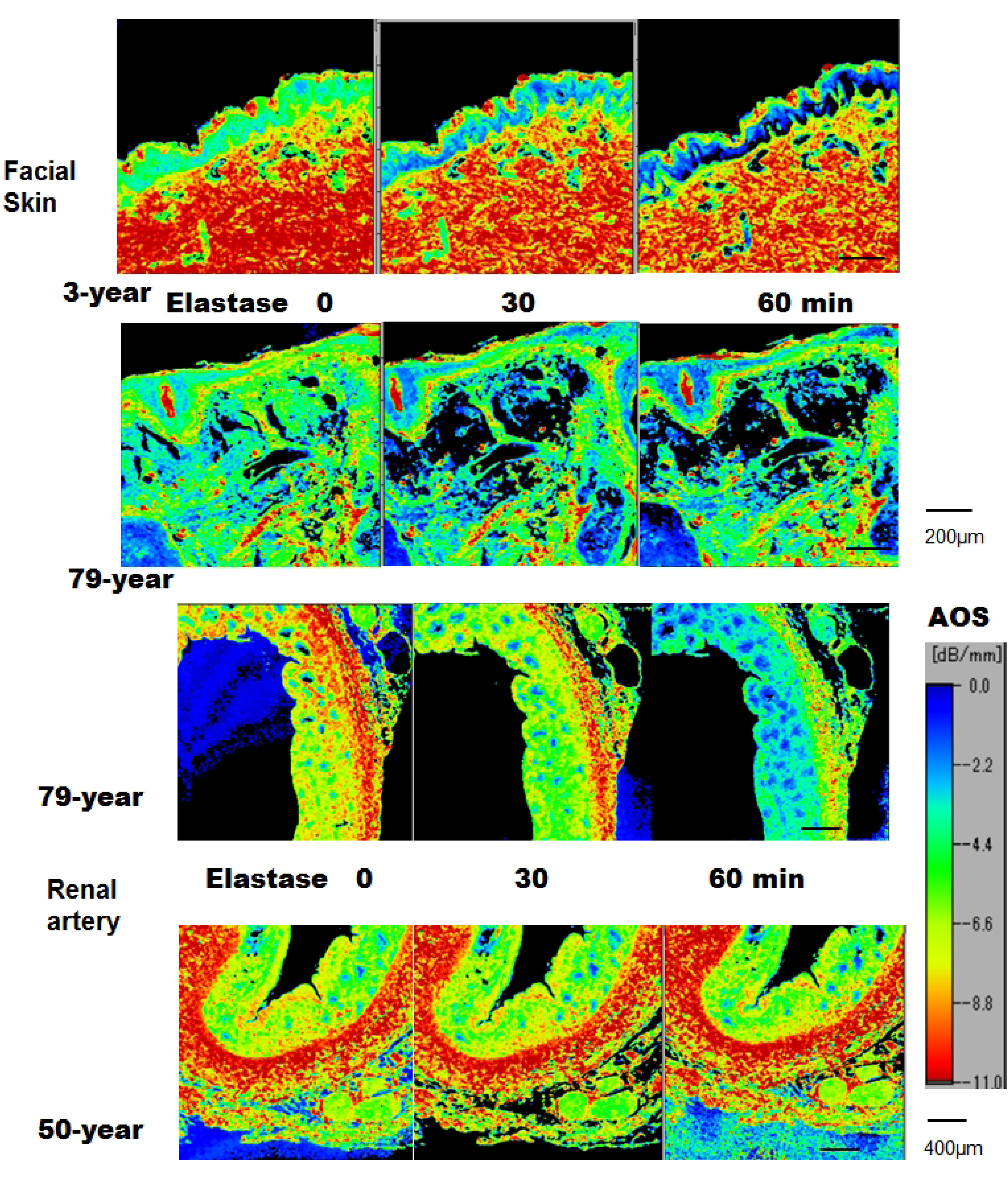
Facial skin from young and elderly individuals and renal arteries digested by elastase. The facial skin sections from young and elderly individuals (upper rows) and renal arteries (lower rows) were digested by elastase. The attenuation-of-sound (AOS) images followed the histology. Regarding the facial skin, the dermis from a young individual displayed high-AOS values and no remarkable changes over time, whereas the dermis from an elderly individual showed reduced AOS values, especially the papillary dermis, which is made up of actinic elastosis. Regarding the epidermis, the epidermis from a young individual promptly exhibited a reduced AOS, whereas the epidermis from an elderly individual gradually declined with some parts maintaining constant values, such as the basal layer. Regarding the renal arteries, the renal arteries from an elderly individual revealed reduced AOS values for all layers, whereas the renal arteries from a young individual maintained constant high-AOS values after digestion.

Fig. 4 shows the corresponding LM pictures with a resorcin stain for elastic fibres before and after digestion. Young skin revealed well-preserved elastic fibres in the dermis, whereas elderly skin showed lost or reduced elastic fibres.

**Fig 4.**
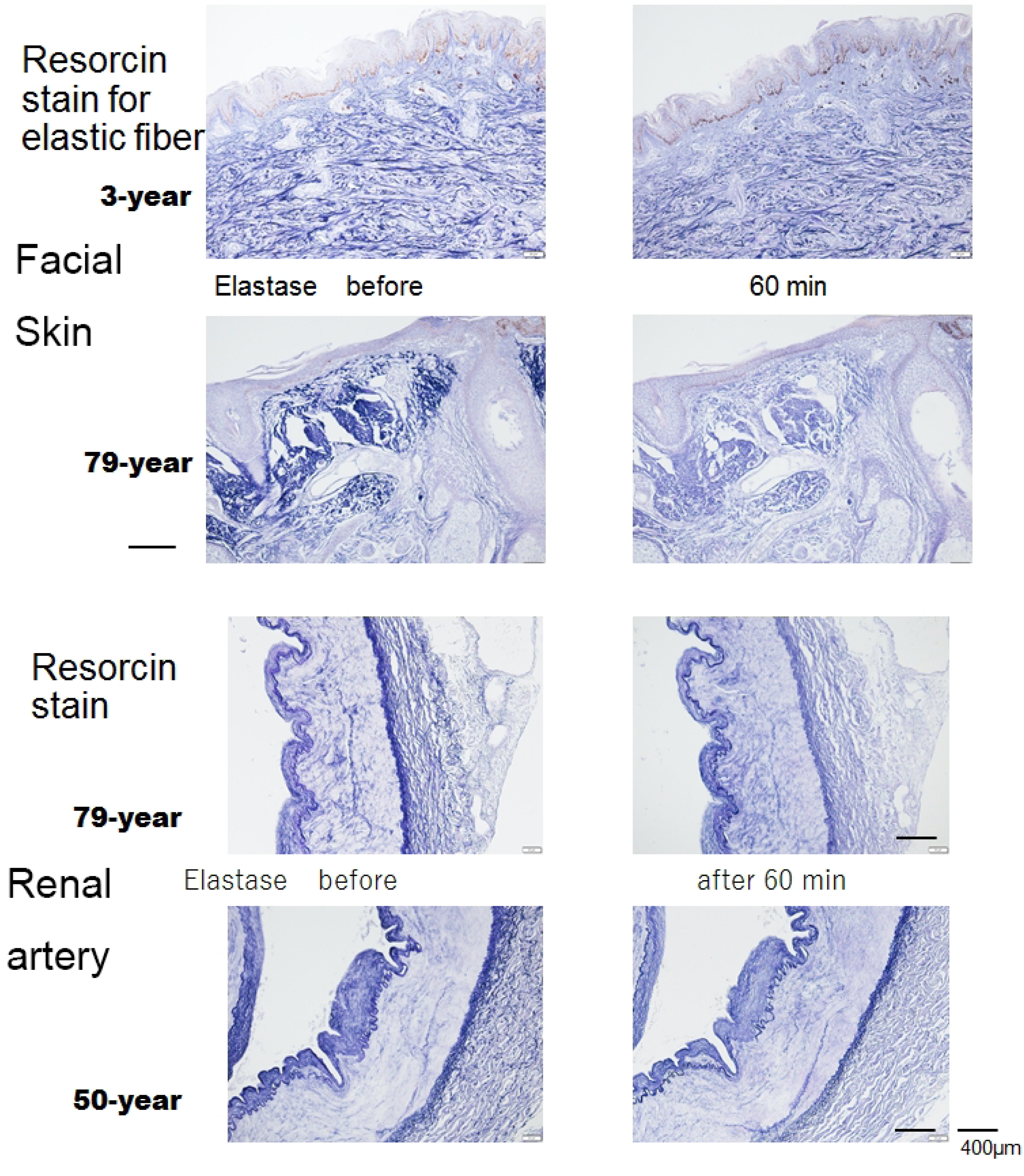
Resorcin staining for elastic fibres before and after skin and renal artery digestion. Regarding the skin, resorcin was bound to the elastic fibres before digestion. After digestion, the dermis of a young individual showed no conspicuous changes, whereas the dermis of an elderly individual waned after digestion. Regarding the epidermis, sections from young and elderly individuals showed no remarkable changes. Regarding the renal artery, the renal artery from an elderly individual showed lost elastic fibres compared with that of a young individual. However, light microscopy histology revealed no remarkable changes.

### Renal arteries of elderly and younger patients digested by elastase

We observed the digestion of renal arteries of elderly and younger individuals by elastase over time. The AOS images (Fig. 3 lower) are compared with the elastic fibre staining before and after digestion (Fig. 4). In the renal artery of young individuals, the elastic fibres of the adventitia included thick and smooth muscles and the elastic fibres of the tunica media were well-developed. In the renal artery of elderly individuals, the elastic fibres of the adventitia were thinner and the tunica medial smooth muscles and elastic fibres were sparse and split.

Elastic fibres in the elderly were easily degraded, showing rapidly decreasing AOS values. On the other hand, the renal artery of young individuals showed little change in the AOS values. In the LM figures, there was a slight decrease in the positive staining of elastic fibres in the elderly and no remarkable changes were observed.

### Digestion of amyloid deposited on the cervical carotid artery with actinase

Amyloid fibrils, which are structurally stable against proteolytic enzymes in general, were digested with actinase over time.

Fig. 5 shows the gradual disappearance of tissue components except amyloid in AOS images. The corresponding images in Congo red staining showed diffuse nonspecific staining before digestion and conspicuous amyloid deposits were retained after digestion.

**Fig 5.**
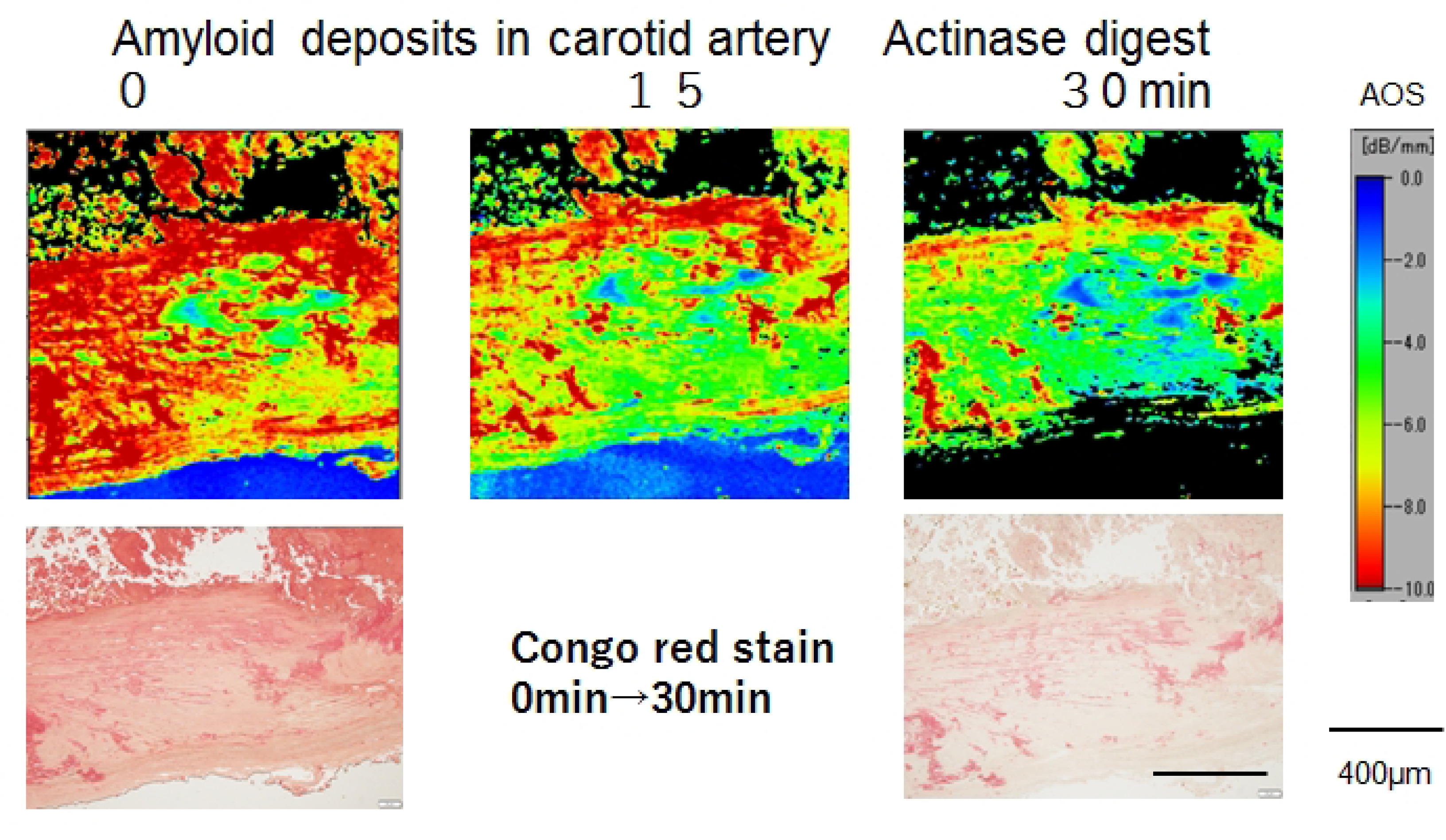
Actinase digestion of amyloid in the carotid artery. The Congo red stain image at the lower left shows broad nonspecific positive areas on the carotid artery before digestion. Figures in the upper row display the attenuation-of-sound (AOS) images after actinase digestion over time. Amyloid, which is resistant to actinase digestion, retained high-AOS values after digestion. Only the amyloid portion remained in the Congo red staining and nonspecific staining disappeared.

### Lymph node with metastatic breast carcinoma digested with actinase

We prepared three different conditions before digestion: (A) untreated, (B) anti-Ki-67 only and (C) anti-Ki-67 with immunostaining using HRP colour development (Fig. 6). After incubation for 1 h with actinase, only section C showed high-AOS values with dots in the lymph node, whereas sections A and B displayed no remarkable changes in the AOS images. The corresponding LM images showed that the positive dots corresponded to the lymphocyte nuclei.

**Fig 6.**
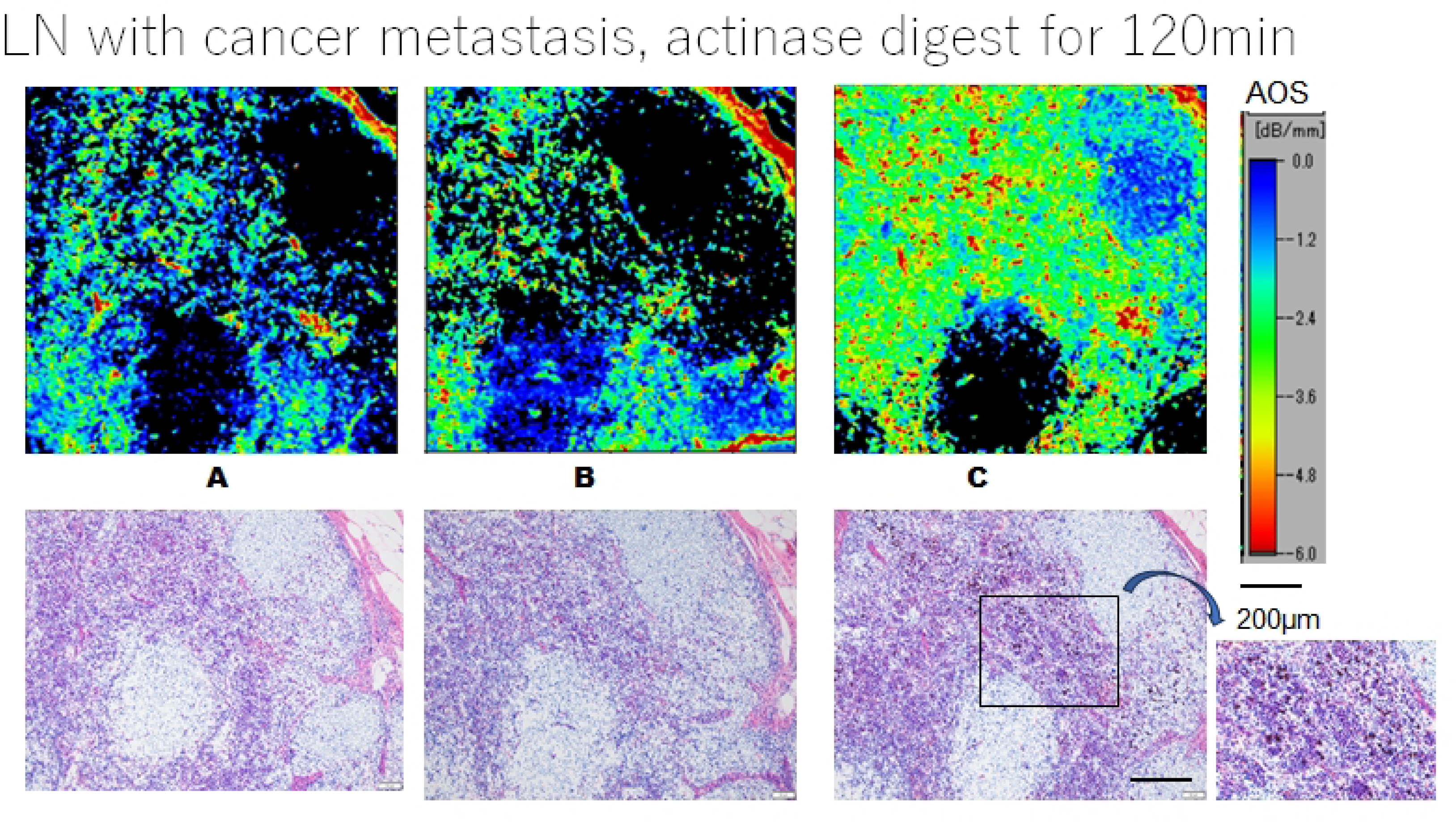
The nucleus was protected from actinase digestion using immunostaining. A lymph node containing metastatic breast carcinoma was digested with actinase for 120 min. Attenuation-of-sound (AOS) images were compared among three different conditions: untreated (A), anti-Ki-67 antibody (B) and anti-Ki-67 antibody with immunostaining reaction using DAB (C). The B and C groups were treated before actinase digestion. A and B slides showed no remarkable changes, whereas slide C exhibited high-AOS-dotted areas corresponding to the immunostained lymphocytes. Figures in the lower row are light microscopy (LM) images that correlate with the above AOS images. Positive brown dots in slice C correspond to the lymphocytes. Upper: AOS image, Lower: LM images before digestion, A: untreated, B: anti-Ki-67 only, C: anti-Ki-67 with immunostaining using DAB

### Corpola amylacea in the brain was digested with amylase

The CA has nothing to do with amyloid; rather, it is derived from sugar-like globules that stain with PAS stain [17,18]. Before digestion, CA was present in tissues surrounding the ventricles (Fig. 7 upper). Digestion with amylase for 1 h revealed that most of the CA surrounding the ventricle disappeared as seen via PAS staining and other CAs scattered in the parenchyma remained. However, the AOS images revealed that most CAs maintained high values even after digestion.

**Fig 7.**
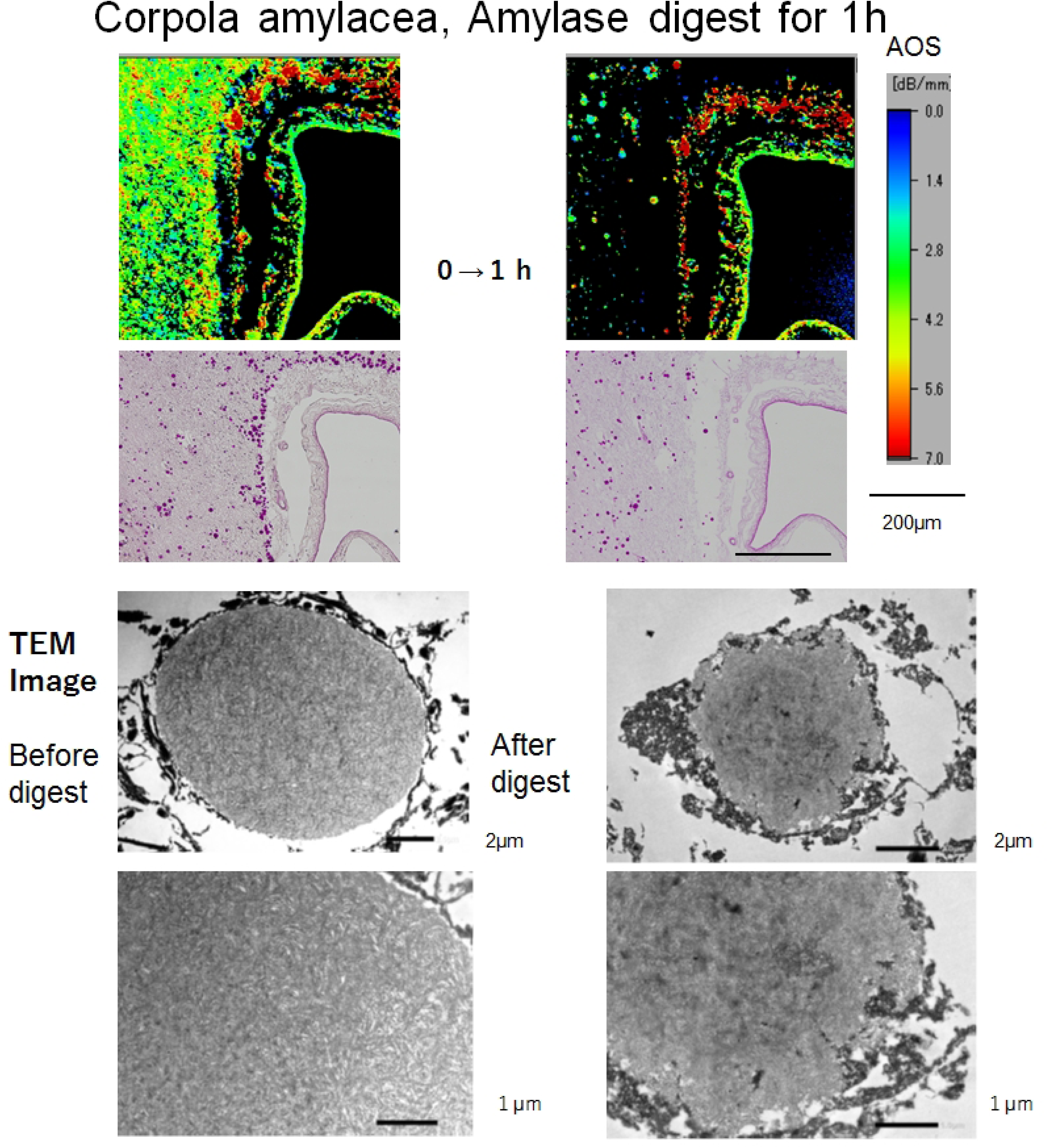
Amylase digestion of the corpora amylacea (CA) of the brain. Upper: attenuation-of-sound (AOS) and PAS-stained images; Lower: transmission electron microscopic (TEM) images A cerebral section from an elderly individual was digested with amylase. The CA surrounding the ventricle disappeared in PAS staining, although AOS images revealed that the CA remained with high-AOS values. Some CA in the parenchyma exhibited a reduced number. Regarding the TEM images, before digestion, the CA was spherical and surrounded by an electron-dense amorphous material. The contents were composed of distorted fibrous structures and electron-dense amorphous material. After digestion, the spheroid was eroded, exhibiting a moth-eaten pattern, which resulted in an irregular shape.

TEM imaging (Fig. 7 lower) showed that the predigested CA was spherical and surrounded by electron-dense amorphous materials. Distorted fibrous structures and electron-dense amorphous material dominated the contents of the spherical body. However, after digestion, the spheroid was eroded with a moth-eaten appearance, which resulted in an irregular shape. The dotted electron-dense materials surrounding the spherical body increased in number.

### DAB reaction products inhibit DNase activity

We examine DAB colour development as an inhibitor of DNase activity. For cytology, a serous adenocarcinoma positive for anti-Ki-67 was digested with DNase for up to 60 min (Fig. 8A). The results showed that 70% of the cancer cells that were larger than the inflammatory cells showed a brown colour in the nucleus in the DAB reaction. In contrast, other negative cells displayed a blue colour in the nucleus by haematoxylin staining before digestion. After 60 min of digestion, the positive cells retained their brown colour, whereas the negative cells diminished in blue or disappeared. We followed the digestion reaction with AOS imaging. In the beginning, most cells showed higher AOS values and then after digestion for 30 min, the AOS values were rapidly reduced, and after 60 min, they decreased further. The separate larger cells or clustered cells remained compared with the smaller cells that were broken in shape.

**Fig 8.**
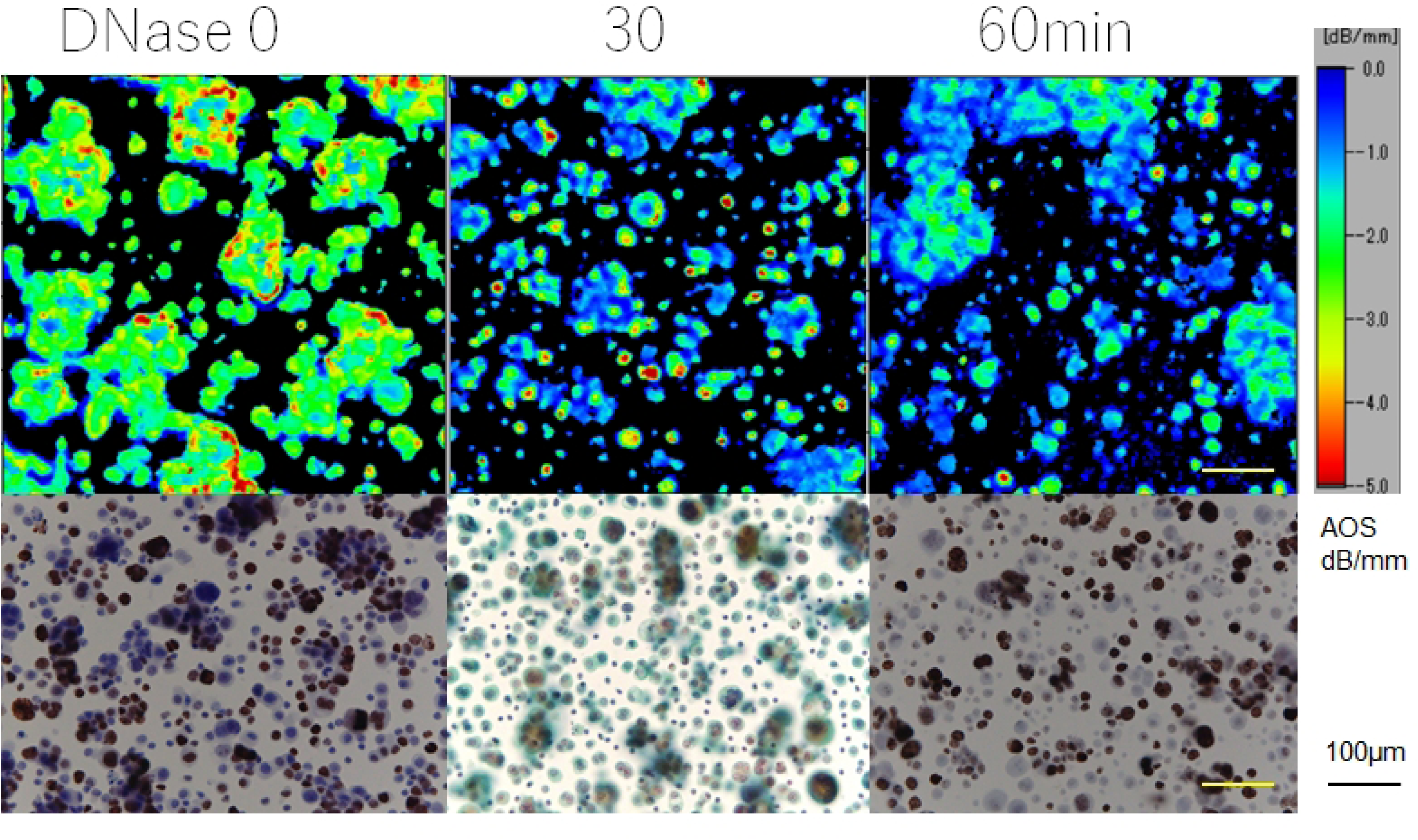
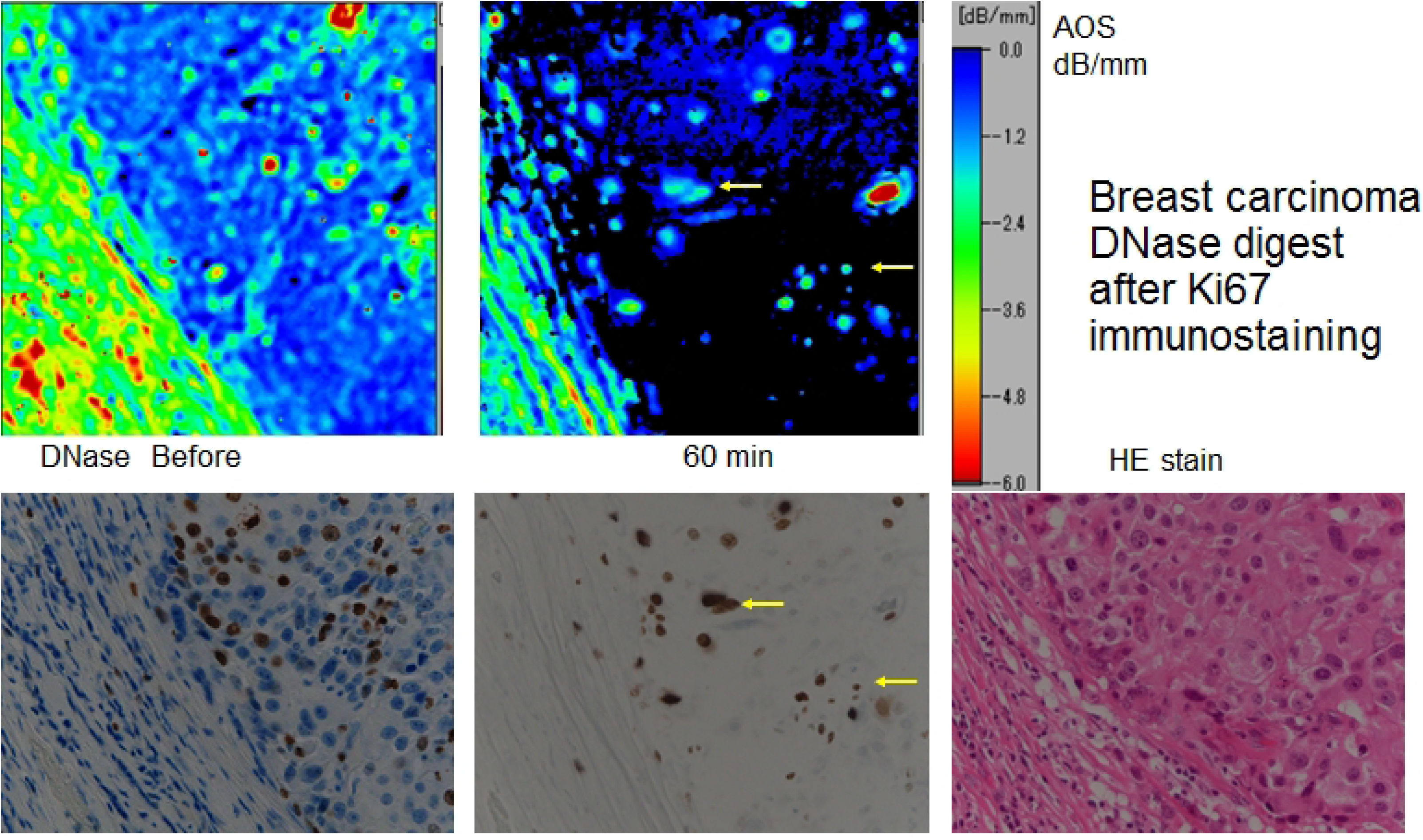
Inhibition against DNase by the DAB reaction products. A. Adenocarcinoma cytology slide digested with DNase after immunostaining The cytological specimen of ascites containing serous carcinoma and inflammatory cells was stained with anti-Ki-67 and digested by DNase. The attenuation-of-sound (AOS) image monitored the reaction at 30 and 60 min of digestion (upper row). All cells presented higher AOS values before digestion and the AOS values gradually decreased, especially in the smaller cells. Separate larger cells or those forming clusters maintained higher AOS values. Light microscopy (LM) images before (lower left) and 60 min after digestion (lower right) showed positive brown spots in the nucleus. Many cancer cells with large nuclei were positive. The nucleus of cells that were stained with haematoxylin before digestion almost disappeared after digestion. Papanicolaou-stained slide is in the centre of the lower row. B. Breast apocrine carcinoma digested by DNase after anti-Ki-67 immunostaining DNase digested the breast carcinoma stained with anti-Ki-67 antibody and DAB solution. The lower row shows the LM slides before and after digestion for 60 min. After digestion, cells stained with haematoxylin disappeared, whereas DAB-positive cells with a brown colour remained. The upper row shows AOS images before and after 60 min of incubation. After digestion, the AOS values of most cells and fibrous tissues declined, except for some cells that correspond to the brown-coloured positive cells (arrows) that maintained high-AOS values.

For the tissue section, breast apocrine carcinoma was digested with DNase after anti-Ki-67 immunostaining (Fig. 8B). After 60 min of digestion, 50% of the cancer cells were positive and maintained their positivity. In contrast, negative cells stained in blue lost their colour after incubation. In the AOS images, cancer cells with high-AOS values were reduced in number after degradation and the residual cancer cells correspond to the positive cells in the LM image.

### Inhibitory effect of haematoxylin on DNase digestion

A section of lymph node with breast carcinoma was digested with DNase. The haematoxylin-stained section was compared with the non-stained section to examine the inhibitory effect of haematoxylin on digestion. The AOS images of the haematoxylin-stained section retained a higher portion of dotted cells compared with the non-stained section (Fig. 9). LM images that were stained with haematoxylin after digestion showed more intense nuclear staining in the pre-stained section than the unstained one. The portion of dotted positive cells with high-AOS values corresponded to the cell nuclei.

**Fig 9.**
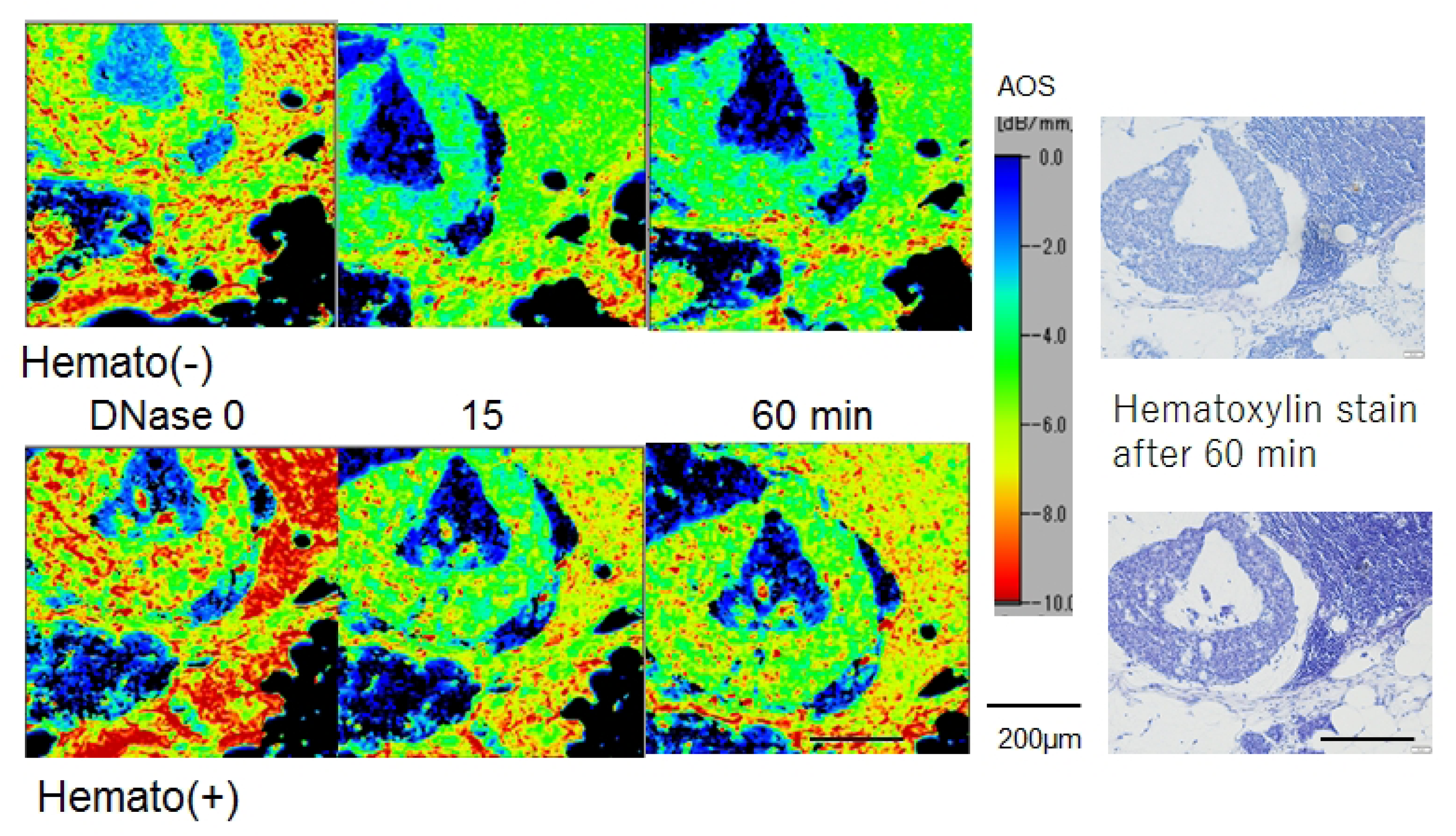
DNase activity was inhibited by haematoxylin in the lymph node with metastatic breast carcinoma. Nuclear digestion by DNase was compared using haematoxylin staining before and after digestion. Attenuation-of-sound (AOS) images with haematoxylin-stained sections showed the nuclear dot structure over time, whereas the unstained section displayed waning nuclear structures. The corresponding light microscopy image after digestion displayed that haematoxylin treatment maintained the nucleus clearer than the untreated section.

## Discussion

### Collagenase type II degradation of the mouse bone

As columns, beams and braces provide support for a house, the trabeculae, cartilage, periosteum and ligaments hold the bones, which mainly consist of collagens. The order of the AOS values corresponded to the fibre density, as seen in Fig. 2. From the manufacturer’s literature, collagenase type 2 is a crude collagenase made from *Clostridium histolyticum* and has high levels of protease activity. The AOS values of the components decreased at almost the same rate. The digital AOS values were easy to compare among the different components and incubation durations.

### Elastase degradation in the facial skin of elderly and young individuals

The facial skin from young and elderly individuals exhibited different reactions to elastase digestion. The dense dermis from young individuals maintained high-AOS values to withstand digestion, whereas the loose dermis from elderly individuals revealed reduced AOS values, especially in the solar elastic portions. The epidermis of young individuals homogeneously showed lower AOS values, while those from elderly individuals were maintained in some areas over time. Although proteins in the skin from elderly individuals included some alterations, such as sun exposure, the degenerated elastic fibres were digestible with elastase. However, the epidermis of elderly individuals revealed focal resistance to elastase digestion.

### Renal arteries from young and elderly individuals digested by elastase

The AOS images revealed conspicuous differences in the renal arteries between young and elderly individuals with elastase digestion. The renal arteries from elderly individuals displayed a remarkable reduction after digestion, which indicates that arteries from elderly individuals were more susceptible to enzyme damage. This explains that the arteries lose elasticity with advancing age. The LM image revealed no remarkable changes between before and after digestion (Fig. 4), which reveals that the AOS image was superior to the LM image following elastase digestion.

### Digestion of amyloid deposited on the cervical carotid artery with actinase

The nonspecific binding of Congo red gradually decreased after actinase digestion, followed by AOS imaging. This method helped to identify tiny amyloid deposits in many contaminants. Moreover, the recovered amyloid from the section is available for analysing the type of amyloid using immunostaining or other molecular techniques.

### Lymph node with metastatic breast carcinoma digested by actinase

Among the three different methods, only the HRP colour development method helped to retain the antigen-positive cells in the section. This antigen-keeping method applies to any antigen that is detected by various antibodies.

### Corpola amylacea in the brain digested with amylase

Although the PAS-stained LM images of the CA disappeared after digestion, the AOS images retained high-intensity values. This discrepancy occurred from the different CA contents. The CA was recently renamed wastesome [19], which consists of sugar chains and proteins, such as immunoglobulins. The sugar chain portion was digested by amylase, while the protein portion remained. The proportion of the sugar chain to that of the protein differed in each CA and the CA showed various AOS values after digestion. The TEM images displayed the fibrillar and electron-dense amorphous contents of the CA. Comparing the TEM images before and after digestion confirmed that amylase digested CA from the surroundings.

### DAB reaction products inhibit DNase activity

Figs 6A and 6B show that DAB-positive cells maintained their positivity after DNase digestion, whereas negative cells disappeared with digestion, which indicates that DAB suppressed the DNase activity. Ki-67-positive cells, which are proliferating cells, can be reserved to analyse DNA sequences. Microdissection methods [20] are used in molecular biology for DNA analysis. This method physically captures cell clusters from the section and requires a special instrument or slides. However, the present method uses routine histology slides and requires no special instruments.

### Inhibitory effect of haematoxylin on DNase digestion

Haematoxylin staining before digestion inhibited DNase activity to maintain the nuclear structure and AOS values compared with the unstained section.

SAM images have the advantage of establishing a numerical evaluation of each structure because the handling data are digital and thus statistical analysis can be performed. Moreover, we could follow the enzymatic degradation process using AOS images over time. The decline in AOS values showed that the enzyme properties could determine the substrate distribution. Furthermore, dyes and antibodies that bind to the substrate could inhibit enzymatic degradation. Therefore, we could intentionally delete or retain the substrate in the section by controlling the enzymatic reaction.

This method has several limitations that should be acknowledged. First, SAM requires flat 10-µm-thick sections, but enzymatic digestive reactions can cause surface irregularities and delamination of the sections, which makes observation challenging. Enzymatic reactions must occur at a suitable pH and temperature and sections are prone to alkali, acid and heat denaturation, which may cause bias in the evaluations. Second, dyes and antibodies bind to tissues and may change the AOS values. Comparisons with untreated and other treated sections are necessary to evaluate the inhibitory effects. Third, determining the observation areas and quality control requires the corresponding LM sections. LM observation is superior in terms of having an adjustable resolution and wider survey area, whereas SAM observation has limited survey areas with a fixed resolution.

## Conclusions

The present method can visualise the location of the substrate in sections and estimate the constituents of complex structures, as seen in CA. Enzymes can discriminate among various substances, such as proteins, sugars and fats. The degradation pattern of the substance differed by age, case and coexisting substance and was comparable with the decline in AOS values.

Furthermore, revealing the binding of some drugs and chemicals to tissues may be possible if changes in AOS values can be observed. The retention of particular cells, nuclei or materials was possible with dyes or antibodies that inhibit enzyme activity. In addition, biochemical or molecular analysis may be available for the residual materials.

We obtained distinct results by degrading various tissues with specific enzymes, and this method can intentionally delete or retain components in the section. Moreover, the degree of degradation can be adjusted and compared using AOS values.

## Acknowledgements

The authors acknowledge Drs. Yuki Egawa and Toshiaki Moriki of Shizuoka City Hospital for the sample preparation, Dr. Yuki Kurita of Department of Preeminent Research Support for making histology sections, and Dr. Isao Ohta of ultrastructure analysis center for making TEM section. Drs. Kanna Yamashita, Michio Fujie, and Toshi Nagata of the Department of Health Science, Hamamatsu University School of Medicine provided the research facilities for this study.

